# A potential role for the gut microbiota in the specialisation of *Drosophila sechellia* to its toxic host noni (*Morinda citrifolia*)

**DOI:** 10.1101/526517

**Authors:** C Heys, AM Fisher, AD Dewhurst, Z Lewis, A Lizé

## Abstract

Adaptation to a novel food source can have significant evolutionary advantages. The fruit fly, *Drosophila sechellia*, is a specialist of the toxic plant noni (*Morinda citrifoli*a). Little is known as to how *D. sechellia* has become resistant to the toxins in the fruit - comprised predominantly of octanoic acid - but to date, the behavioural preferences for the fruit and genetic architecture underlying them, have been well studied. Here, we examine whether the gut microbiota could have played a role in adaptation to the fruit. In the first series of experiments, we examine the gut microbiota of wild-type, laboratory reared flies and characterise the gut microbiota when reared on the natural host plant, versus a standard *Drosophila* diet. We show a rapid transition in the core bacterial diversity and abundance within this species and discover sole precedence of *Lactobacillus plantarum* when reared on *M. citrifolia*. We also discover that flies reared on a laboratory diet are more likely to carry bacterial pathogens such as *Bacillus cereus*, although their function in *Drosophila* is unknown. Flies reared on a laboratory diet have a significantly reduced weight but with no impact on the risk of death before adulthood, when compared to the wild noni diet. In the second series of experiments, we examine the potential role of the gut microbiota in adaptation to octanoic acid resistance in this species and its sister species, *Drosophila melanogaster*, to which the fruit is usually fatal. We use a combination of methods to analyse resistance to octanoic acid by conducting life history analysis, behavioural assays and bacterial analysis in both *D. sechellia* and *D. melanogaster*. We find that by creating experimental evolution lines of *D. melanogaster* supplemented with gut microbiota from *D. sechellia*, we can decrease *D. melanogaster* aversion to octanoic acid, with the flies even preferring to feed on food supplemented with the acid. We suggest this represents the first step in the evolutionary and ecological specialisation of *D. sechellia* to its toxic host plant, and that the gut microbiota, *Lactobacillus plantarum* in particular, may have played a key role in host specialisation.

## Introduction

Many studies have examined the complex relationships between animals and plants (reviewed in Herrera and Pellmyr, 2009). The dynamic ecological and evolutionary interactions between an animal and its host plant can take many forms. In some cases, insects have even become adapted to living on a toxic plant host, such as *Pierinae* butterflies that feed on toxic *Brassicales* (e.g. Edger et al., 2015). Developing resistance to a toxic host plant through specialisation can have a multitude of advantages, including the ability to exploit an otherwise unutilised resource, and there are many ways that insects can overcome the toxins in the host plant. Behavioural and physiological adaptations to exploit toxic resources can drive speciation of insects or other animals (Matsuo et al., 2007). However, it is argued that specialisation to a toxic host does not necessarily result in speciation itself, and therefore the role of ecological specialisation in speciation remains to be proven (Matsuo et al., 2007).

The way in which these specialists adapt to life in a novel environment can occur through a number of different ways, and one is potentially through the gut microbiota (e.g. Bolnick et al., 2014; Morrow et al., 2015). In insects, for example, members of the order Hemiptera have evolved to feed on plant phloem sap - a nutritionally poor diet due to the grossly unbalanced amino acid composition (e.g. Douglas, 1993; Sandström and Moran, 1999; Sandström, 2000). A number of studies have demonstrated that all phloem feeders within this order possess certain symbiotic bacteria that mitigate the effects of this nutritionally poor diet (Buchner 1965; Gündüz and Douglas, 2009). For example, two specialist species of Lepidoptera, *Hyles euphorbiae* and *Brithys crini*, feed exclusively on latex-rich *Euphorbia* sp. and alkaloid-rich *Pancratium maritimum*, respectively (Vilanova et al., 2016). Metagenomic sequencing has identified that the primary microbiota within the gut is Entereococcus sp., which it is predicted to be responsible for mitigating the effects of these nutrient-poor diets to the host.

The participation of the gut microbiota in the transformation of toxic plant chemicals is an important aspect to be considered when studying insect-plant interactions, and one that is gaining attention (e.g. Genta et al., 2006; Ceja-Navarro et al., 2015). Yet due to the complexity and diversity of gut microorganisms, there is little experimental evidence to support this idea. It was suggested by Douglas (1992) that a possible role of the midgut microbiota is in detoxification of toxic compounds and a study by Genta et al., (2006) in *Tenebrio molitor* highlighted the role of gut microbiota in detoxifying the cell walls of fungi and bacteria that typically inhabit their food source. Similarly, the coffee berry borer (*Hypothenemus hampei*), the most devastating insect pest of coffee crops worldwide, has adapted to metabolise caffeine – a toxic alkaloid present within coffee plants (Ceja-Navarro et al., 2015). Caffeine is shown to be degraded in the gut via *Pseudomonas* species, which subsist on caffeine as a sole source of carbon and nitrogen (Ceja-Navarro et al., 2015).

The majority of species within the *Drosophila melanogaster* species-complex are food generalists and saprophagous, meaning they feed on a variety of decaying plant matter (Rohlfs and Kürschner, 2010). Studies have shown the importance of a diverse diet in creating and maintaining a diverse gut microbiota within *D. melanogaster*, with a diverse gut microbiota increasing survival and reducing development time (Rohlfs and Kürschner, 2010). In comparison, several species within the *D. melanogaster* species-complex have evolved some form of diet specialisation (e.g. Lachaise et al., 1988). One species within this group, *Drosophila sechellia*, is a specialist of ripe *Morinda citrifolia* fruit – a toxic fruit commonly known as the Tahitian noni (Jones, 2005). *D. sechellia’s* closely related species, *Drosophila simulans, Drosophila mauritiana* and *D. melanogaster* are notably repelled by the pungent scent of the fruit and even die upon contact (Legal et al., 1994; Legal et al., 1999). Resistance of *D. melanogaster* to the fruit is shown to be dependent on strain (Legal et al., 1992).

The primary toxins found within noni are octanoic and hexanoic acids (Farine et al., 1996), and numerous studies have focussed on the underlying genetic and molecular mechanisms that have enabled *D. sechellia* to adapt to the fruit (e.g. Harada et al., 2008, 2012; Labeur et al., 2002; Matsuo et al., 2007. However, whether the gut microbiota has played a role in this specialisation has not yet been investigated. Interestingly, Chandler *et al.* (2011) characterised the microbiota of wild *D. sechellia* found feeding on the noni and discovered that the gut is dominated by a single *Lactobacillales* OTU (84%). This demonstrates the very low bacterial community richness and diversity within this species, particularly when it is compared to its sister species, *D. melanogaster*, which exhibit greater diversity and carry the bacterial genera *Lactobacillus*, *Acetobacter* and *Enterococcus* (Ryu et al., 2008; Brummel et al., 2004; Ren et al., 2007; Cox and Gilmore, 2007). Further, the composition of an individual’s gut microbiota is known to be influenced by pH (see Overend et al., 2016). Overend et al. (2016) demonstrated that decreasing the pH in certain regions of the *Drosophila* gut can lead to an increased abundance of key members of the gut microbiota – *Lactobacillus* and *Acetobacter*. This raises the question whether the very low bacterial richness found in the *D. sechellia* gut when it is feeding on its acidic natural host diet, *M. citrifolia*, is due to the pH determining the microbiota? Alternatively, potentially the almost exclusive prevalence of this *Lactobacillales* is caused by the specialism of *D. sechellia* to the host plant, and thus the *Lactobacillales* acts as a form of detoxifying agent by metabolising the toxic acids found within the noni.

In this study we investigated the role of the gut microbiota on host specialisation in *D. sechellia*. This study is separated into two sections. In the first, we determined the effect that rearing *D. sechellia* on a standard laboratory diet has on the diversity and richness of the gut microbiota. *D. sechellia* are widely kept in the laboratory, but little attention has been paid to the effect that feeding this specialist species a generalist diet has on the resulting gut microbiota. Flies were first reared on a standard *Drosophila* diet (ASG), then moved onto noni, before being transferred back onto ASG. At each stage, the diversity and abundance of the gut bacteria was measured. We also disentangled the role of pH on shaping the gut microbiota, from the toxic compounds present in the noni. We introduced a new dietary treatment, salak fruit (*Salacca zalacca*), with similar nutritional and acidic properties to noni but lacking in the toxic compounds - octanoic and hexanoic acid. We determined the effect that these diets - a standard laboratory diet, noni and salak fruit - had on a series of life history traits, including larval, pupal and adult weight and the risk of death before adulthood. We predicted that the gut microbiota would become more simplified on the noni diet, but not the salak or ASG, which would indicate that the gut microbiota plays a role in specialisation to this diet.

In the second part of this experiment, we investigated the effects of differing concentrations of the toxic compound octanoic acid, the main acidic constituent of *D. sechellia’*s natural host plant, *M. citrifolia*, on weight, development time, survival, bacterial load and diversity in *D. sechellia*. We also investigated the same effects in *D. sechellia’s* sister species, *D. melanogaster*, to determine any differences between a fruit specialist and a fruit generalist species. We analysed both inbred and outbred lines in *D. melanogaster* in order to test the effect of strain on ability to withstand octanoic acid exposure. We predicted that *D. melanogaster* would have a reduced weight, development time and survival ability compared to *D. sechellia*, particularly in the inbred strain in which genetic diversity is low. We also predicted that we would observe differences in the diversity and abundance of the gut microbiota of *D. sechellia* compared to *D. melanogaster*, due to *D. sechellia* possessing a more specialised gut microbiota that enable them to withstand high concentrations of octanoic acid. We further examined the role of the gut microbiota in this specialisation by creating experimental evolution lines of *D. melanogaster* supplemented with *D. sechellia* gut microbes. As we hypothesised the gut microbiota may have played a key role in the specialisation in *D. sechellia*, we predicted that after ten generations *D. melanogaster* would be less averse to exposure to octanoic acid.

## Materials and methods

### Flies used and general maintenance

*D. sechellia* stocks were obtained from the National Drosophila Species Stock Center located in San Diego. Three lines of outbred flies were utilised (lines 0.21, 0.07 and 0.08), that were collected on Cousin Island, Seychelles in 1980 and maintained in the laboratory ever since. All flies were kept and reared at 25°C on a 12:12hour light-dark cycle. Flies were kept in standard 75×25mm *Drosophila* vials containing 25ml of standard *Drosophila* food composed of yeast/agar/maize/sugar. Flies were moved to new vials every 4 days.

### Experiment 1: The changing gut microbiota of *D. sechellia*

#### Experimental treatments

Newly emerged, virgin adult flies were obtained from the stock population and transferred to a new vial containing 25ml of a standard *Drosophila* dietary media composed of yeast/sugar/agar/maize (hereon known as ASG). Flies were left to mature on this media for two days before being transferred to a new vial containing the same media (N=30). After one week, two males and two females from different vials from each stock line were isolated using carbon dioxide gas anaesthesia. The protocol below detailing the bacterial analysis was then followed for these individuals, to determine the gut bacterial load and diversity of flies reared on this diet. These flies formed the “ASG 1” treatment.

The remaining flies were then gently aspirated into fresh vials containing 25g of noni and left for one week (N=30). After this time, two males and two females from different vials from each stock line were again isolated and the same bacterial analysis protocol was followed. This enabled us to determine any changes in the gut microbiota in the same population of flies, that were first reared on a different diet. These flies formed the “Noni” treatment.

Similarly, the remaining flies were again, gently aspirated into fresh vials containing 25ml of ASG and left for one week (N=30). After this time, two males and two females from different vials from each stock line were again isolated and the same bacterial analysis protocol was followed. This enabled us to determine any further changes in the gut microbiota, when flies from the same population were transferred between two different diets in a short period of time. These flies formed the “ASG 2” treatment.

#### pH and diet type on life history traits

In order to test the effect of acidity on *D. sechellia* life history traits, we constructed three different diets of varying pH. Flies were reared on one of three diets: ASG (for 1l of water: 85g of sugar, 60g of corn, 20g of yeast, 10g of agar and 25ml of nipagin), noni, or salak fruit. *Morinda citrifolia* is the diet of wild *D. sechellia* and has a low, acidic pH of 3.86 due to the high concentrations of both octanoic and hexanoic acids (Legal et al., 1994). The salak fruit diet was used as an alternative diet to the noni, as it also has a low, acidic pH at 3.59, but does not contain the toxic octanoic and hexanoic acids that are present in noni. These two diet types were compared to the typical laboratory diet of ASG, that has a higher, more alkaline pH than the fruit diets at 5.97.

#### Risk of death before adulthood

The number of days was measured from day of female oviposition to day of adult emergence. Vials were checked at three time points within each day – 9am, 12pm and 5pm – and the cumulative number of adults emerged from each time point was scored (N_ASG_ = 76; N_Noni_ = 60; N_Salak_ = 125).

#### Weight at different life stages

In order to accurately determine the effect of acidity on life history, the weights of three different life stages were measured. For larval weight, vials were checked daily during the morning and any third instar larvae present were removed and washed with distilled water in order to remove any excess food. Larvae were grouped according to treatment and placed into the freezer at −18°C for two hours. Later, larvae were removed and weighed using an Ohaus five place balance and their weight was recorded (in mg) to four decimal places (N_ASG_ = 50; N_Noni_ = 50; N_Salak_ = 50).

For the pupal and adult weights, vials were similarly checked daily at three time points – 9am, 12pm and 5pm – to check for any freshly pupated or newly emerged individuals. For the pupae, care was taken to remove pupae from the vials without damaging them (N_ASG_ = 50; N_Noni_ = 50; N_Salak_ = 50). The adult flies were isolated as virgins and separated according to sex. Adults were placed into vials at a standard density of ten per vial and left for two hours to allow their wings to dry out and inflate. Two hours later, vials were placed into the freezer at −18°C and left overnight. Pupae were grouped according to treatment and placed into the freezer at −18°C for two hours. Later, the pupae and adults were removed and weighed using an Ohaus five place balance and their weight was recorded (in mg) to four decimal places. In the adults, male and female measurements for each treatment were recorded and analysed separately (females: N_ASG_ = 53; N_Noni_ = 50; N_Salak_ = 50; males: N_ASG_ = 45; N_Noni_ = 45; N_Salak_ = 50). This method of determining adult weight was used for both sections of the experiments.

### Experiment 2: Octanoic acid resistance in *D. sechellia* and *D. melanogaster*

#### Survival rate

To determine resistance to the toxic octanoic acid compound, the survival of the three different strains, outbred *D. melanogaster*, inbred *D. melanogaster* and *D. sechellia*, to exposure of differing concentrations of octanoic acid was measured. Newly emerged, virgin adults were isolated from the stock populations and transferred to fresh vials containing 25ml standard *Drosophila* media and left for 3 days to mature. During this time, ≥99% octanoic acid (purchased from Sigma-Aldrich) was diluted in distilled water to the following concentrations: 0% (distilled water), 1%, 5%, 10%, 25%, 50%. 30μl of the acid solute was pipetted onto the surface of a fresh vial containing 25ml standard *Drosophila* media. The vial was tipped to the side to ensure the acid covered the entire surface of the food media and left to dry for 2 hours. After this time, males and females were separated according to sex and placed at a standard density of 10 individuals per vial. The number of dead individuals was counted every 24hours for a total period of 168hours and the survival rate calculated (N_InbredDmel_ = 50; N_OutbredDmel_ = 50; N_Dsechellia_ = 50 for each acid concentration).

#### Development time

Mated adults at a density of five females and five males were placed onto fresh vials containing the following concentrations of octanoic acid: 0% (distilled water), 1%, 5%, 10%, 25%, 50%, to determine the effects of different concentrations of octanoic acid on development time. The pairs were left to oviposit. Development time was measured as the number of days from female oviposition to day of adult emergence. Vials were checked at three time points within each day – 9am, 12pm and 5pm – and the cumulative number of adults emerged from each time point was scored. Emergent adult flies from each time point were removed from the vial and placed into a fresh vial containing 25ml of standard *Drosophila* food (sample sizes are documented below.

#### Experimental evolution of *D. melanogaster* lines exposed to *D. sechellia* gut microbiota

To determine whether it is the gut microbiota that enables *D. sechellia* to become attracted to and feed on a diet that contains high levels of the toxic compound, octanoic acid, we created a series of experimental evolution lines. Using the stock population of outbred *D. melanogaster*, we added *D. sechellia* gut solute to the dietary media over several generations. Here, stock population *D. sechellia* that were continually reared on a noni diet, were first surface sterilised in 70% ethanol, rinsed in distilled water and air dried. The head was then removed. Two guts were dissected into each Eppendorf containing 250μl of sterile LB (Lysogeny Broth) broth. An equal number of males and females were used to ensure there were no sex-specific differences in the bacterial content. Gut tissue was homogenised with a sterile plastic pestle. 30μl of gut isolate was then pipetted onto the surface of a vial containing 25ml standard *Drosophila* media. The vial was tipped to the side to ensure the solute covered the entire surface of the food media and left to dry for 20minutes. After this time, newly emerged, virgin males and females were isolated from the stock population of outbred *D. melanogaster* and placed at a standard density of 10 males and 10 females per vial. After pupae were seen in all vials, the adult flies were removed so the offspring and adults did not interbreed. Once the offspring had emerged as adults, a sub-sample of the first generation were isolated. Some of these individuals were harvested and their gut bacterial content and diversity analysed. The rest of the sub-sample of the first generation were placed into octanoic acid aversion trials (see protocol below). The remaining flies were placed onto a fresh vial similarly containing 25ml standard *Drosophila* media and 30μl of gut isolate. This process was repeated until the offspring reached the 10^th^ generation. Newly emerged adult offspring were harvested as before. The remaining adults were placed into octanoic acid aversion trials. Thus, the aversion trials were conducted on unselected stock individuals for each species, and the first and tenth generations of the experimental evolution lines.

#### Octanoic acid aversion trials

In order to test whether experimental evolution of outbred *D. melanogaster* reared on a diet supplemented with *D. sechellia* gut microbiota reduces aversion to the toxic octanoic acid, aversion trials were performed, using a similar methodology to that utilised previously (e.g. Dekker et al., 2006). Newly emerged, virgin adults were isolated from the *D. sechellia* and outbred *D. melanogaster* stock populations, separated according to sex and placed into fresh vials containing 25ml standard *Drosophila* media. Similarly, newly emerged, virgin adults were isolated from the first generation (hereon known as Dmel 1) and 10^th^ generation (hereon known as Dmel 10) experimental evolution lines described above, separated according to sex and placed into fresh vials containing 25ml standard *Drosophila* media. Flies were left to mature for 3 days before being placed into an aversion arena (Figure 1). The aversion arena consisted of a standard petri dish (measuring 100mm × 15mm) containing two pieces (10g) of standard *Drosophila* food located at opposite ends, with a marked line half-way across clearly showing the two separate sides. 10ul of concentrated octanoic acid was pipetted on to one of these pieces of food. An individual fly was gently aspirated into the centre of the arena and left to acclimatise for five minutes. After this time, the side on which the fly was located was scored as its preference, i.e., either the side with food containing octanoic acid, or without (female N_Dmel_ = 50, male N_Dmel_ = 50; female N_Dsech_ = 50, male N_Dsech_ = 50; female N_Dmel1_ = 51, male N_Dmel1_ = 50; female N_Dmel10_ = 49, N_Dmel10_ = 48).

**Figure 1.**
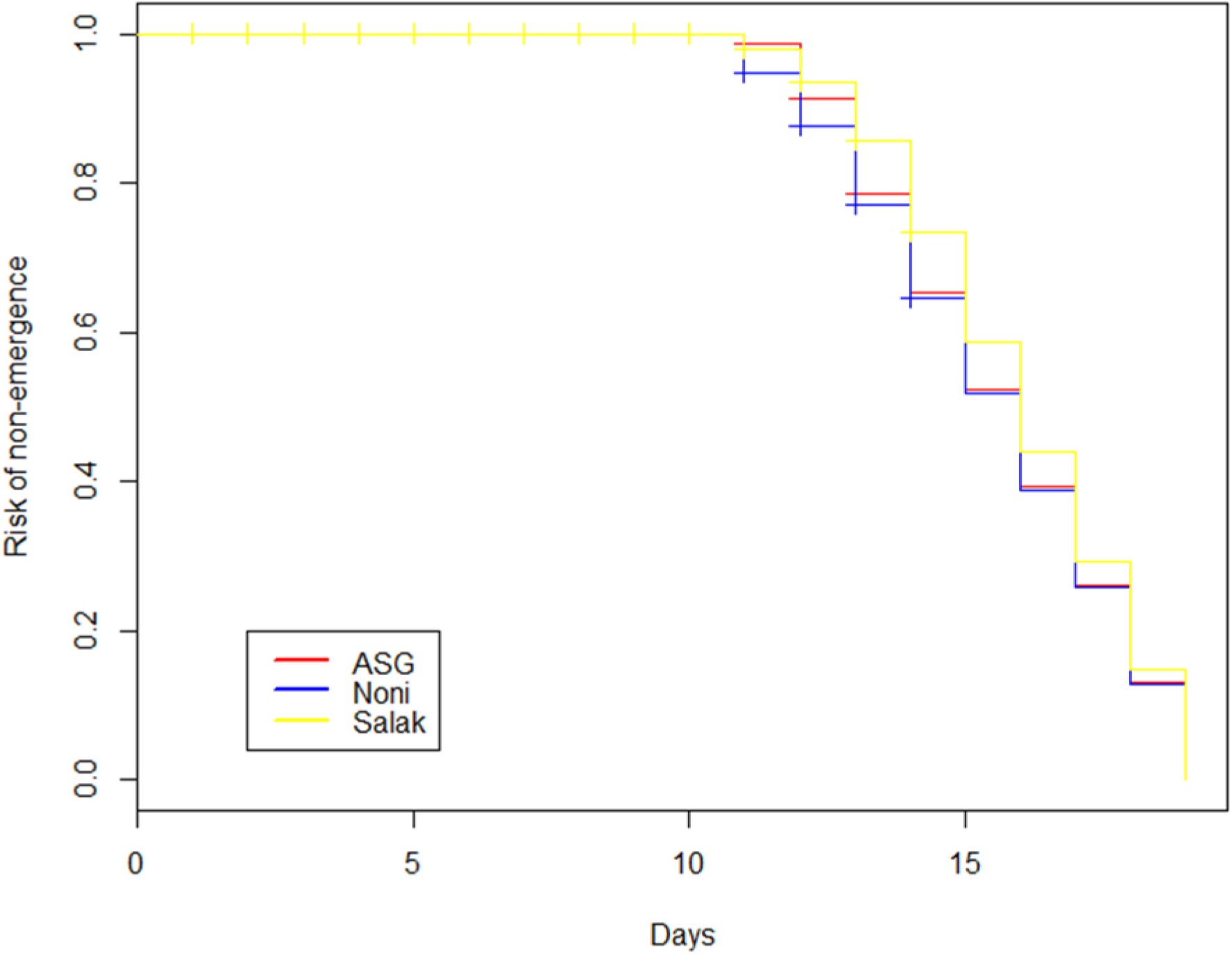
Development time failure, measured in days as the risk to die before adulthood, of *D. sechellia*. Eggs were reared under one of three different dietary treatments - either ASG, noni or salak diets.

#### Bacterial analysis for all experiments

Collected flies were first surface sterilised in 70% ethanol, rinsed in distilled water and air dried. The head was then removed. Two guts were dissected into each Eppendorf containing 250μl of sterile LB (Lysogeny Broth) broth (Bertani, 2004). An equal number of males and females were used to ensure there were no sex-specific differences in the bacterial content. Gut tissue was homogenised with a sterile plastic pestle. 100μl of gut homogenate was pipetted onto BHI (Brain, Heart Infusion) agar (Atlas, 2004) and spread-plated using a sterile glass loop. BHI media was used as it was found to favour greater colony growth. Plates were left to air dry aseptically, before being closed and sealed with parafilm. Plates were incubated at 25°C for 72 hours, and bacterial load was quantified by performing CFU (Colony Forming Unit) counts.

Single colonies were isolated using a sterile 1μl loop and placed into an Eppendorf with 10μl sterile water. PCR amplification was performed in a 25μl reaction volume consisting of 10μl nuclease-free water, 13μl Taq green master mix, 0.5μl of forward primer 27F (5’-AGAGTTTGATCMTGGCTCAG-3’) and reverse primer 1492R (5′-GGTTACCTTGTTACGACTT-3’) and 1μl of template DNA. Thermal cycling was performed for 90 seconds at 95°C as initial denaturation, followed by 35 cycles of 30 sec at 95°C for denaturation, 30 sec at 55 °C as annealing, 90 sec at 72 °C for extension, and final extension at 72 °C for 5 min. 1500 bp 16S PCR products were purified with Ampure beads and subjected to Sanger sequencing. The resulting sequences were identified using NCBI BLAST against the nt database (Altschul et al., 1990).

### Statistical analysis

All data were analysed using R (version 3.3.0; R Core Team, 2016). Larval, pupal and adult weight were analysed using separate General Linear Models (GLM). Adult weight was first analysed with both sexes grouped, before further analysis separated according to sex were performed. The aversion data was analysed using a binomial GLM. Variation in development time, survival response to octanoic acid and the risk of death before adulthood was analysed via Cox Proportional-Hazard Regressions.

Development failure of flies was used as the ‘event’ for the risk of death before adulthood data. The *Survdiff* function was used to assess differences between two or more survival curves according to treatment. The *coxph* function was used to assess differences between treatments. This allowed treatments to be compared in a pairwise fashion, to ascertain whether all treatments differed, or whether any significant differences observed were derived from a single treatment.

## Results

### Experiment 1: The changing gut microbiota of *D. sechellia*

Bacterial colony growth was observed in all treatments, with both greater diversity and greater abundance of bacteria found in the ASG 1 and ASG 2 flies (Table 1). Flies analysed from these treatments were found to have *Lactobacillus plantarum*, *Paenibacillus sp*. and *Bacillus cereus* species present. In nearly all of the noni flies only *L. plantarum* was observed, with the exception of minor colony growth in two of the male replicates. Little difference was observed between the three different strains of *D. sechellia*, or between sexes.

**Table 1.** Number of bacterial colonies isolated from the midgut of adult flies. Flies were first reared on ASG (represented by ASG 1), then moved onto noni, before being transferred back onto ASG (ASG 2).

| **Diet** | **Strain** | **Replicate** | **Sex** | ***L. plantarum*** | ***Paenibacillus* sp.** | ***Bacillus cereus*** |
| --- | --- | --- | --- | --- | --- | --- |
| ASG 1 | 0.21 | 1 | F | 3.12x10<sup>2</sup> | 0.20x10<sup>1</sup> | 0 |
| ASG 1 | 0.21 | 2 | F | 1.82x10<sup>2</sup> | 1.40x10<sup>1</sup> | 0 |
| ASG 1 | 0.21 | 1 | M | 1.58x10<sup>2</sup> | 0 | 0.10x10<sup>1</sup> |
| ASG 1 | 0.21 | 2 | M | 2.50x10<sup>1</sup> | 0.90x10<sup>1</sup> | 0 |
| ASG 1 | 0.07 | 1 | F | 5.11x10<sup>3</sup> | 1.81x10<sup>2</sup> | 0.50x10<sup>1</sup> |
| ASG 1 | 0.07 | 2 | F | 5.94x10<sup>3</sup> | 1.23x10<sup>2</sup> | 0.20x10<sup>1</sup> |
| ASG 1 | 0.07 | 1 | M | 4.88x10<sup>3</sup> | 5.40x10<sup>1</sup> | 0.70x10<sup>1</sup> |
| ASG 1 | 0.07 | 2 | M | 3.58x10<sup>3</sup> | 1.75x10<sup>1</sup> | 2.20x10<sup>1</sup> |
| ASG 1 | 0.08 | 1 | F | 6.25x10<sup>3</sup> | 2.02x10<sup>2</sup> | 2.70x10<sup>1</sup> |
| ASG 1 | 0.08 | 2 | F | 4.09x10<sup>3</sup> | 1.96x10<sup>2</sup> | 1.50x10<sup>1</sup> |
| ASG 1 | 0.08 | 1 | M | 3.17x10<sup>3</sup> | 2.70x10<sup>1</sup> | 0.40x10<sup>1</sup> |
| ASG 1 | 0.08 | 2 | M | 2.89x10<sup>3</sup> | 8.80x10<sup>1</sup> | 0 |
| Noni | 0.21 | 1 | F | 2.72x10<sup>3</sup> | 0 | 0 |
| Noni | 0.21 | 2 | F | 1.78x10<sup>3</sup> | 0 | 0 |
| Noni | 0.21 | 1 | M | 1.62x10<sup>2</sup> | 0 | 0 |
| Noni | 0.21 | 2 | M | 1.59x10<sup>2</sup> | 0 | 0 |
| Noni | 0.07 | 1 | F | 1.43x10<sup>3</sup> | 0 | 0 |
| Noni | 0.07 | 2 | F | 1.34x10<sup>3</sup> | 0 | 0 |
| Noni | 0.07 | 1 | M | 2.55x10<sup>2</sup> | 1.50x10<sup>1</sup> | 0 |
| Noni | 0.07 | 2 | M | 1.92x10<sup>2</sup> | 0.70x10<sup>1</sup> | 0 |
| Noni | 0.08 | 1 | F | 4.5x10<sup>2</sup> | 0 | 0 |
| Noni | 0.08 | 2 | F | 1.81x10<sup>2</sup> | 0 | 0 |
| Noni | 0.08 | 1 | M | 8.10x10<sup>1</sup> | 0.50x10<sup>1</sup> | 0 |
| Noni | 0.08 | 2 | M | 7.20x10<sup>1</sup> | 0 | 0 |
| ASG 2 | 0.21 | 1 | F | 1.21x10<sup>3</sup> | 2.40x10<sup>1</sup> | 0 |
| ASG 2 | 0.21 | 2 | F | 1.45x10<sup>3</sup> | 1.20x10<sup>1</sup> | 0.10x10<sup>1</sup> |
| ASG 2 | 0.21 | 1 | M | 1.22x10<sup>2</sup> | 0.20x10<sup>1</sup> | 0.10x10<sup>1</sup> |
| ASG 2 | 0.21 | 2 | M | 2.31x10<sup>2</sup> | 0.50x10<sup>1</sup> | 0.20x10<sup>1</sup> |
| ASG 2 | 0.07 | 1 | F | 4.51x10<sup>2</sup> | 1.50x10<sup>2</sup> | 0.70x10<sup>1</sup> |
| ASG 2 | 0.07 | 2 | F | 5.22x10<sup>2</sup> | 8.90x10<sup>2</sup> | 0.20x10<sup>1</sup> |
| ASG 2 | 0.07 | 1 | M | 2.09x10<sup>3</sup> | 2.90x10<sup>1</sup> | 0.60x10<sup>1</sup> |
| ASG 2 | 0.07 | 2 | M | 2.87x10<sup>3</sup> | 4.60x10<sup>1</sup> | 1.30x10<sup>1</sup> |
| ASG 2 | 0.08 | 1 | F | 3.22x10<sup>3</sup> | 1.98x10<sup>2</sup> | 2.90x10<sup>1</sup> |
| ASG 2 | 0.08 | 2 | F | 2.46x10<sup>3</sup> | 2.51x10<sup>2</sup> | 2.20x10<sup>1</sup> |
| ASG 2 | 0.08 | 1 | M | 1.78x10<sup>3</sup> | 1.12x10<sup>2</sup> | 0.50x10<sup>1</sup> |
| ASG 2 | 0.08 | 2 | M | 2.34x10<sup>3</sup> | 1.43x10<sup>2</sup> | 0.70x10<sup>1</sup> |

#### Risk of death before adulthood

No significant difference was observed in the risk of death before adulthood between the ASG or noni treatments (Z_2_=0.373, P=0.709) (Figure 1). However, flies reared on salak had a significantly lower risk of death before adulthood than flies reared on noni (Z_2_=-2.187, P=0.028). A trend was observed in the risk of death before adulthood between ASG and salak treatments, with salak individuals exhibiting a higher risk of death before adulthood than ASG flies (Z_2_=-1.905 P=0.056).

#### Larval weight

No difference was observed in larval weight in any pairwise comparisons across all three treatments: ASG and noni (T_2_=0.850, P=0.397), ASG and salak (T_2_=0.335, P=0.738), noni and salak (T_2_=-0.515, P=0.608) (Figure 2).

**Figure 2.**
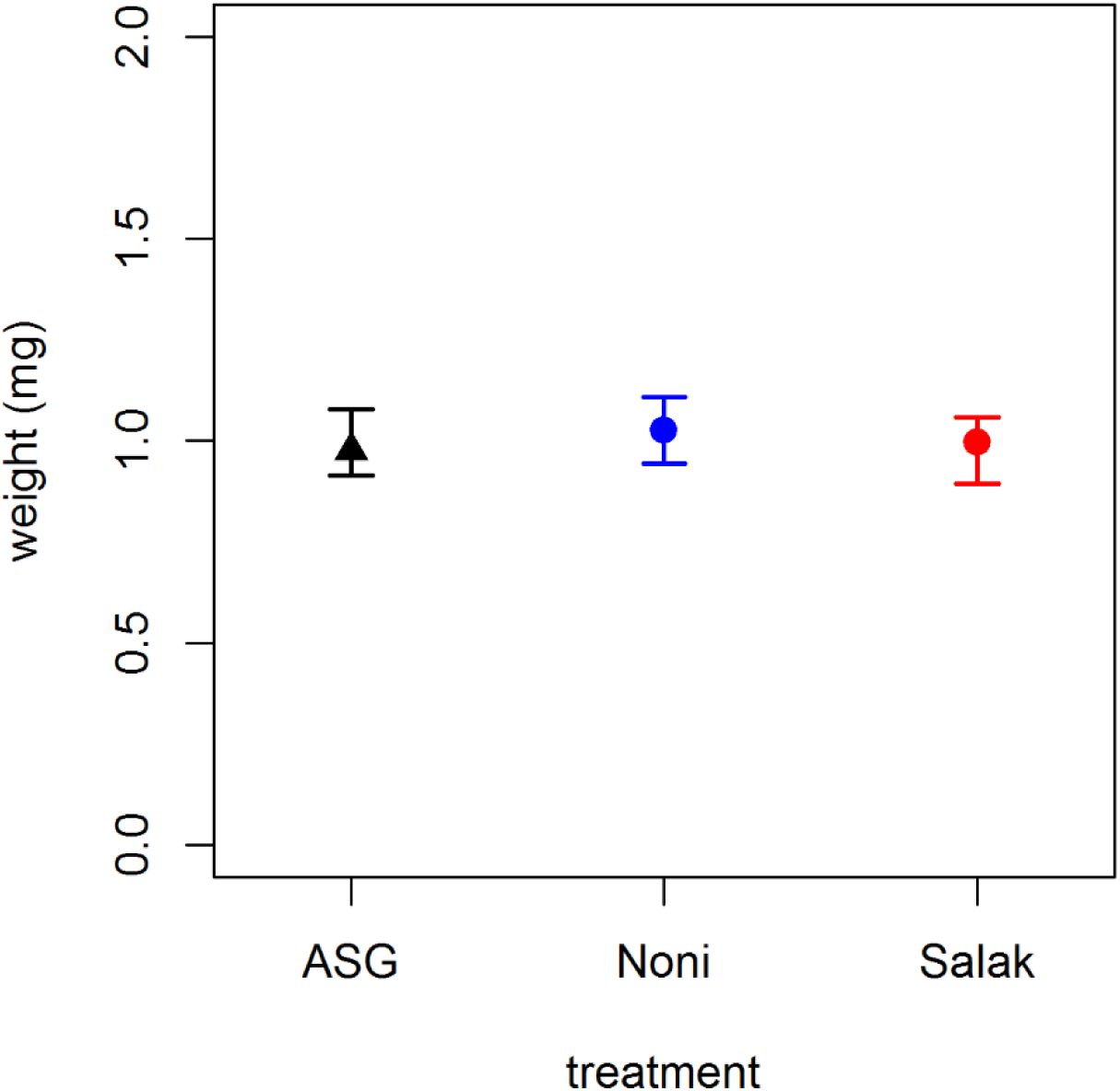
Weight (mg) of third instar *D. sechellia* larvae reared on one of three different diet types - either ASG, noni or salak. No significant differences were found.

#### Pupal weight

Pupae collected from the noni treatment weighed significantly less than pupae from the ASG treatment (T_2_=-8.961, P<0.001) (Figure 3). Similarly, pupae obtained from the salak treatment were found to weigh significantly less than the pupae from the ASG treatment (T_2_=-8.722, P<0.001). No difference in pupal weight was observed between pupae from the noni and salak treatments (T_2_=0.239, P=0.812).

**Figure 3.**
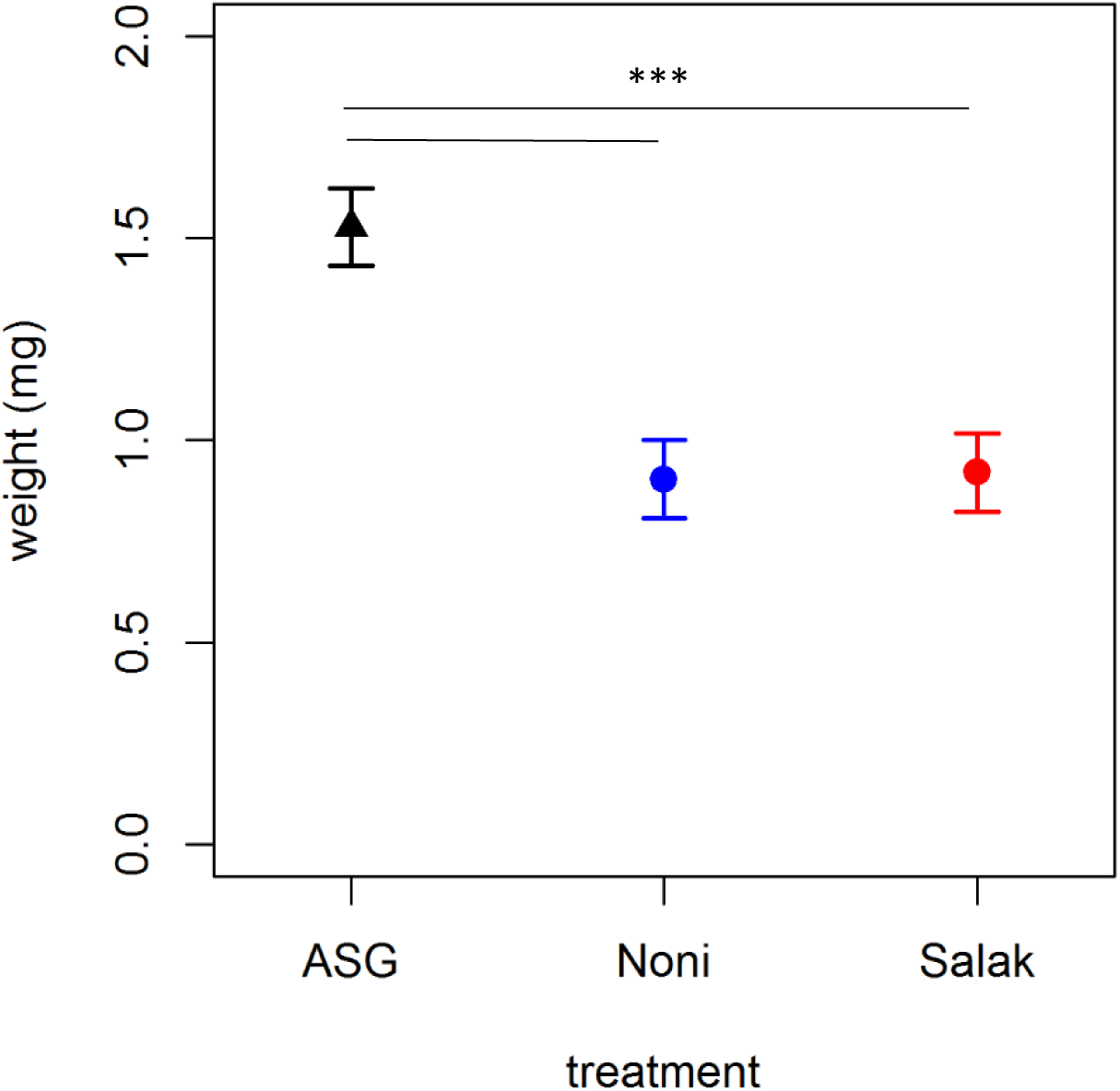
Weights (mg) of *D. sechellia* pupae reared on one of three different dietary treatments - either ASG, noni or salak. Vials were checked at various time points for freshly pupated flies. Significant results are shown with *.

#### Adult weight

Adult male flies were always found to weigh less than females (P<0.001). When males and females are analysed separately, in male flies, significant differences were found across two comparisons, with both noni males and salak males weighing significantly more than ASG flies (T_2_=3.919, P<0.001; T_2_=3.115, P=0.002, respectively) (Figure 4). However, no differences were found between the weights of noni and salak males (T_2_=0.905, P=0.366). In females, significant differences were found across all comparisons, with noni females weighing more than ASG females (T_2_=19.069, P<0.001) and salak females (T_2_=-15.410, P<0.001). Similarly, salak females were also shown to weigh significantly more than ASG females (T_2_=3.435, P<0.001).

**Figure 4.**
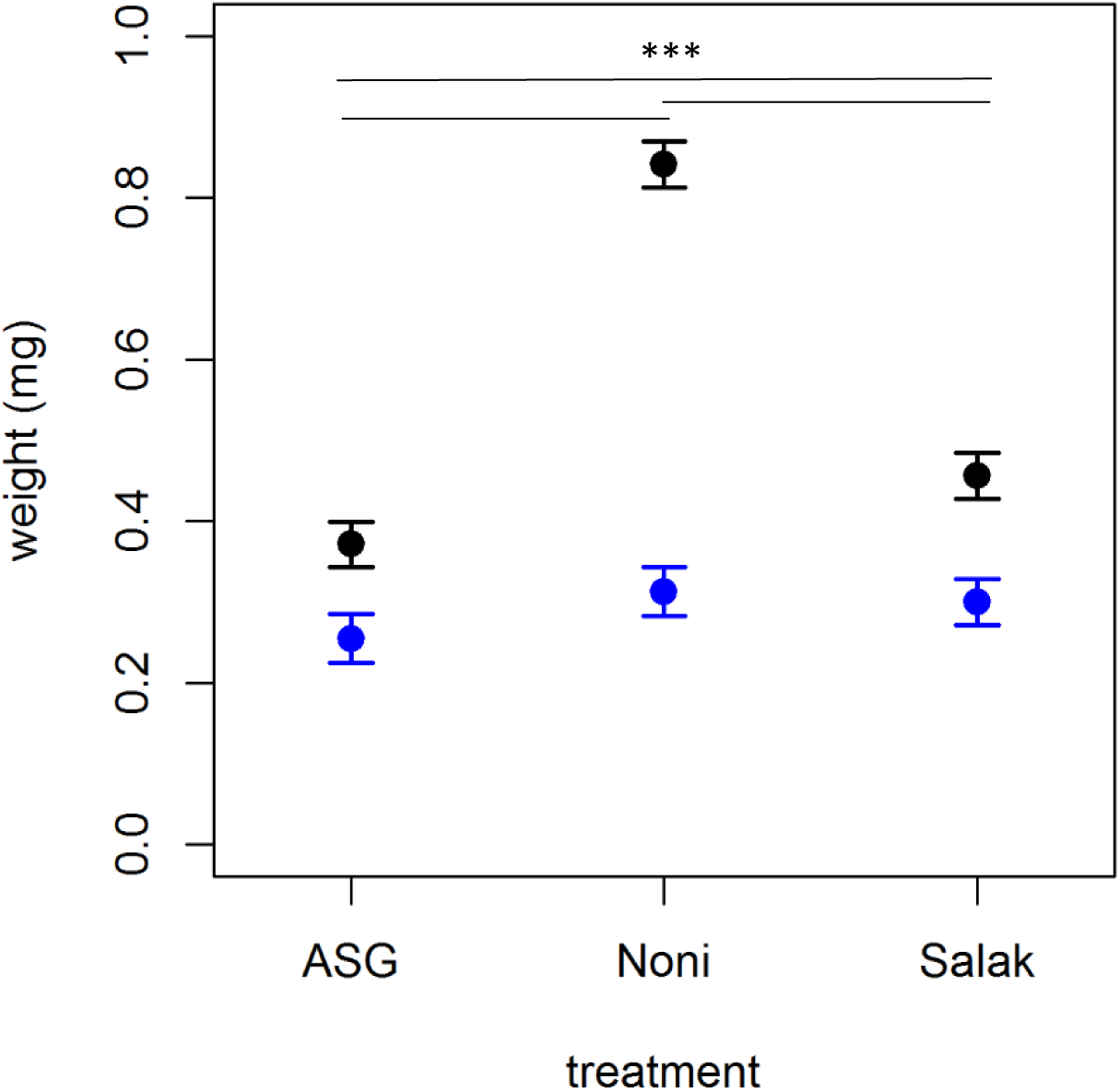
Weights (mg) of both male and female, newly emerged adult *D. sechellia* flies reared on one of three different diet types - either ASG, noni or salak. Newly emerged adults were collected at various time points and allowed two hours for wing inflation. Male flies are shown here using the blue plots, whilst females are depicted in the black plots. Significant results are marked with a *.

#### Bacterial analysis

Bacterial colony growth was observed in all treatments, with a greater diversity of bacteria found in the flies that were reared on the ASG diet (Table 2). In all flies that were reared on the noni and salak treatments, only one kind of bacterial colony formed. Sanger sequencing data identified this as *L. plantarum.* In the ASG diet flies, *L. plantarum* was similarly observed, with *Paenibacillus sp*. and *Bacillus cereus* also found. Little difference was observed between the bacterial load of noni and salak reared flies, or between the different sexes.

**Table 2.** Number of bacterial colonies isolated from the midgut of adult flies that were reared on one of three diets – ASG, noni or salak. Males and females were quantified separately.

| Diet | Replicate | Sex | *L. plantarum* | *Paenibacillus*<br>*sp.* | *B. cereus* |
| --- | --- | --- | --- | --- | --- |
| ASG | 1 | F | 2.52x10<sup>2</sup> | 1.30x10<sup>1</sup> | 0.20x10<sup>1</sup> |
|  | 2 | F | 2.91x10<sup>2</sup> | 1.90x10<sup>1</sup> | 0 |
|  | 1 | M | 1.89x10<sup>2</sup> | 2.30x10<sup>1</sup> | 0 |
|  | 2 | M | 2.34x10<sup>2</sup> | 1.80x10<sup>1</sup> | 0.60x10<sup>1</sup> |
| Noni | 1 | F | 2.85x10<sup>2</sup> | 0 | 0 |
|  | 2 | F | 3.19x10<sup>2</sup> | 0 | 0 |
|  | 1 | M | 1.51x10<sup>2</sup> | 0 | 0 |
|  | 2 | M | 1.99x10<sup>2</sup> | 0 | 0 |
| Salak | 1 | F | 1.31x10<sup>2</sup> | 0 | 0 |
|  | 2 | F | 1.66x10<sup>2</sup> | 0 | 0 |
|  | 1 | M | 2.12x10<sup>2</sup> | 0 | 0 |
|  | 2 | M | 2.45x10<sup>2</sup> | 0 | 0 |

### Experiment 2: Octanoic acid resistance in *D. sechellia* and *D. melanogaster*

#### Survival rate

*D. sechellia* females reared on 0% octanoic acid exhibited significantly higher survival than females reared on the 1% concentration (Z_5_=3.340; P<0.001), the 5% concentration (Z_5_=3.936; P<0.001), the 25% treatment (Z_5_=4.740; P<0.001), or the 50% treatment (Z_5_=4.464; P<0.001) (Figure 5). However, no significant difference was observed between 0% and 10% treatments (Z_5_=1.664; P=0.096). Flies reared on the 10% treatment exhibited significantly higher survival than flies reared on the 25% treatment (Z_5_=3.564; P<0.001) or the 50% (Z_5_=3.448; P<0.001).

**Figure 5.**
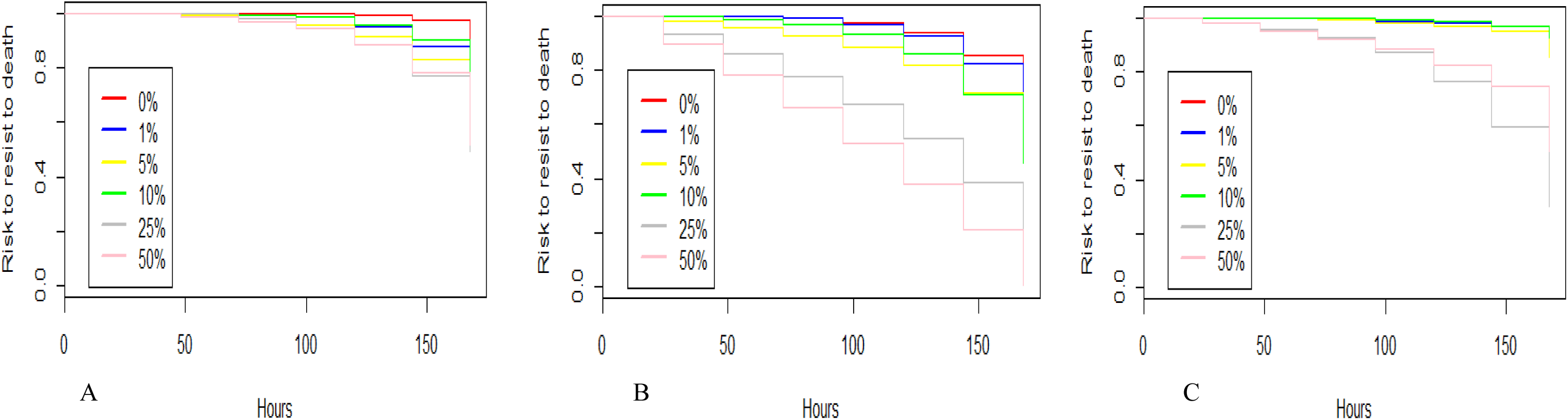
the proportion of survived female flies when exposed to standard *Drosophila* food containing differing concentrations of octanoic acid – 0%, 1%, 5%, 10%, 25%, 50%. The vials were checked every 24 hours with the last time point 168 hours, and the number of dead flies recorded. The graphs are arranged according to species: *D. sechellia* (A), inbred *D. melanogaster* (B) and outbred *D. melanogaster* (C).

In female inbred *D. melanogaster*, no difference was observed in the survival ability of 0% treated and 1% treated flies (Figure 5) (Z_5_=0.778; P=0.436), but 0% treated flies were significantly more able to survive than those reared on 5% (Z_5_=4.020; P<0.001), 10% (Z_5_=3.538; P<0.001), 25% (Z_5_=9.943; P<0.001), or 50% (Z_5_=13.075; P<0.001). Flies reared on the 1% treatment were significantly more able to survive than those at higher concentrations: 5% (Z_5_=3.340; P<0.001), 10% (Z_5_=2.836; P=0.004), 25% (Z_5_=9.689; P<0.001), or 50% (Z_5_=13.102; P<0.001). Females reared at the 50% octanoic acid concentration exhibited significantly higher mortality than those reared at 25% (Z_5_=5.919; P<0.001).

In female outbred *D. melanogaster*, flies reared at a 25% concentration had a significantly higher mortality rate than flies reared on any other treatment: 0% (Z_5_=-7.205; P<0.001), 1% (Z_5_=-7.203; P<0.001), 5% (Z_5_=-7.236; P<0.001), 10% (Z_5_=-6.943; P<0.001), including the 50% concentration (Z_5_=-2.789; P=0.005). Similarly, flies reared at 50% concentration had a significantly higher mortality rate than flies reared at 0% (Z_5_=-5.825; P<0.001), 1% (Z_5_=-5.822; P<0.001), 5% (Z_5_=-5.590; P<0.001) or 10% (Z_5_=-5.816; P<0.001).

When comparing the proportion of survived females across species, there is little difference in survival rates in *D. sechellia* across all octanoic acid concentrations, compared to inbred *D. melanogaster* (Figure 2). Here, females reared at higher concentrations exhibit a higher mortality, with all individuals recorded as dead in the 25% and 50% treatments, unlike in *D. sechellia* in which approximately half the sample were still alive at the end of the trial. The proportion of survived individuals is around 0.3 in outbred *D. melanogaster* at higher concentrations (25% and 50%), with individuals reared at low concentrations (0%, 1%, 5% and 10%) exhibiting similar survival rates to *D. sechellia*.

In contrast to female survival rates, in male *D. sechellia*, flies reared on the 0% acid treatment had equal survival rates to all other treatments, with no difference found compared to 1% reared flies (Z_5_=0.941; P=0.357), 5% (Z_5_=1.341; P=0.180), 10% (Z_5_=-1.448; P=0.148), 25% (Z_5_=0.808; P=0.419) or 50% (Z_5_=-0.045; P=0.964) (Figure 6). Similar to the female *D. sechellia*, male flies reared on the 10% treatment were significantly more able to survive than those reared on 1% treatment (Z_5_=-2.239; P=0.019), the 5% treatment (Z_5_=-2.691; P=0.007) and the 25% treatment (Z_5_=2.207; P=0.027).

**Figure 6.**
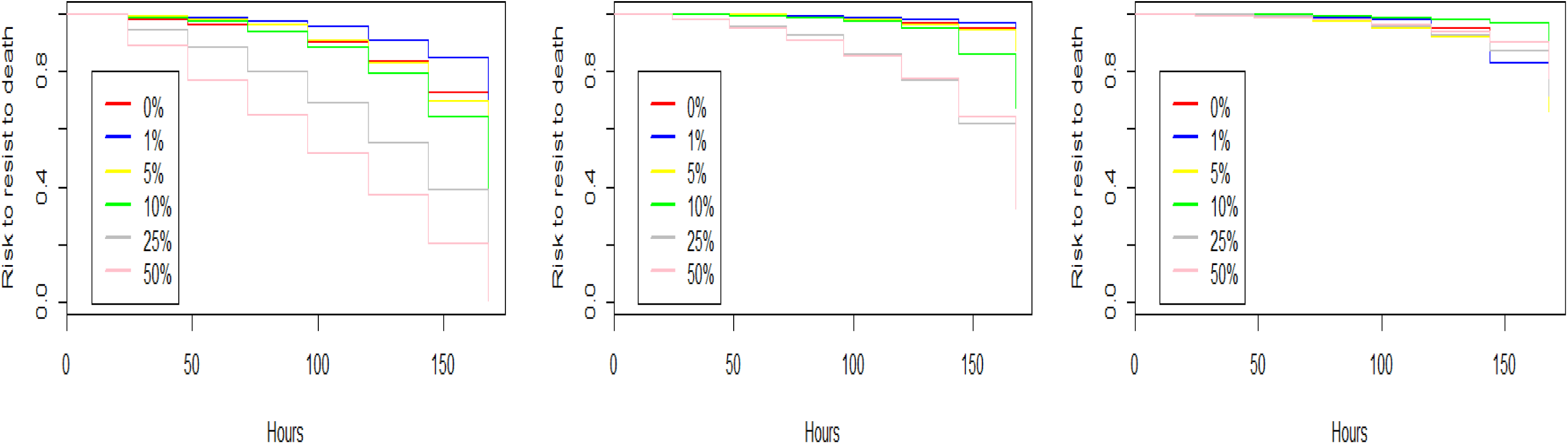
the proportion of survived male flies when exposed to standard *Drosophila* food containing differing concentrations of octanoic acid – 0%, 1%, 5%, 10%, 25%, 50%. The vials were checked every 24 hours with the last time point 168 hours, and the number of dead flies recorded. The graphs are arranged according to species: *D. sechellia* (A), inbred *D. melanogaster* (B) and outbred *D. melanogaster* (C).

In male inbred *D. melanogaster* flies, 0% reared flies had a significantly higher survival ability than 25% flies (Z_5_=7.830; P<0.001) and 50% flies (Z_5_=12.660; P<0.001). Flies reared on the 1% acid concentration had a higher proportion survived than all other concentrations, including 0%: (Z_5_=3.013; P=0.002), 5% (Z_5_=3.405; P<0.001), 10% (Z_5_=4.668; P<0.001), 25% (Z_5_=9.556; P<0.001) and 50% (Z_5_=13.244; P<0.001). A significantly higher mortality was observed in the 25% and 50% treatments compared to both the 5% (Z_5_=7.481; P<0.001, Z_5_=12.430; P<0.001, respectively) and the 10% treatment (Z_5_=6.311; P<0.001, Z_5_=11.670; P<0.001, respectively).

In male outbred *D. melanogaster*, no significant difference in survival ability is found between flies reared at the lower concentrations. Flies reared at 10% concentration had a significantly higher mortality than those reared at 0% (Z_5_=-3.120; P=0.001), 1% (Z_5_=-3.095; P=0.001) and 5% (Z_5_=-2.618; P=0.008) but this was significantly lower than those reared at 25% and 50% (Z_5_=5.770; P<0.001; Z_5_=5.790; P<0.001, respectively). Similarly, flies reared at 25% and 50% had a significantly higher mortality than those reared at 0% (Z_5_=-7.061; P<0.001, Z_5_=-7.074; P<0.001), 1% (Z_5_=-7.035; P<0.001, Z_5_=-7.048; P<0.001) and 5% (Z_5_=-7.026; P<0.001, Z_5_=-7.040; P<0.001). All other pairwise comparisons for survival analysis are non-significant.

When comparing the proportions of survived males across species, there is a vast difference in the final proportion of survived *D. sechellia* males compared to inbred *D. melanogaster* (Figure 6). Male inbred *D. melanogaster* are less able to survive than *D. sechellia* males, particularly when reared at high concentrations (e.g. 25% and 50%). The survival ability of outbred *D. melanogaster* flies is better than that of inbred flies with a higher proportion survived at the end of the trial, even when males were placed at high concentrations, but the proportion was less than that of *D. sechellia.*

#### Development time

In *D. sechellia*, no significant differences were found in the time taken to develop from egg to adulthood between any of the acid treatments (Figure 4). In particular, no significant difference was observed between the 0% and the 50% treatments (F_5_=0.740; P=0.459). However, in the inbred line of *D. melanogaster*, significant differences were observed between all treatments with flies reared on higher concentrations of acid taking longer to develop than those reared on smaller concentrations (Figure 7). Similarly, variation was seen in the time taken for outbred *D. melanogaster* individuals to develop when reared on different concentrations of acid (Figure 4). Flies reared on 0% acid concentration took significantly less time to develop than those reared on 1% treatment (F_5_=-7.930; P<0.001), 5% treatment (F_5_=-4.647; P<0.001), 10% treatment (F_5_=-8.679; P<0.001), 25% (F_5_=-7.808; P<0.001) or 50% (F_5_=-4.900; P<0.001). Similar to the inbred *D. melanogaster*, no significant differences were observed between 10% and 25% reared flies (F_5_=0.759; P=0.447),10% and 50% flies (F_5_=1.733; P=0.083), 25% and 50% flies (F_5_=1.098; P=0.272). All other pairwise comparisons are non-significant. A full summary of all pairwise comparisons is provided in the appendix.

**Figure 7.**
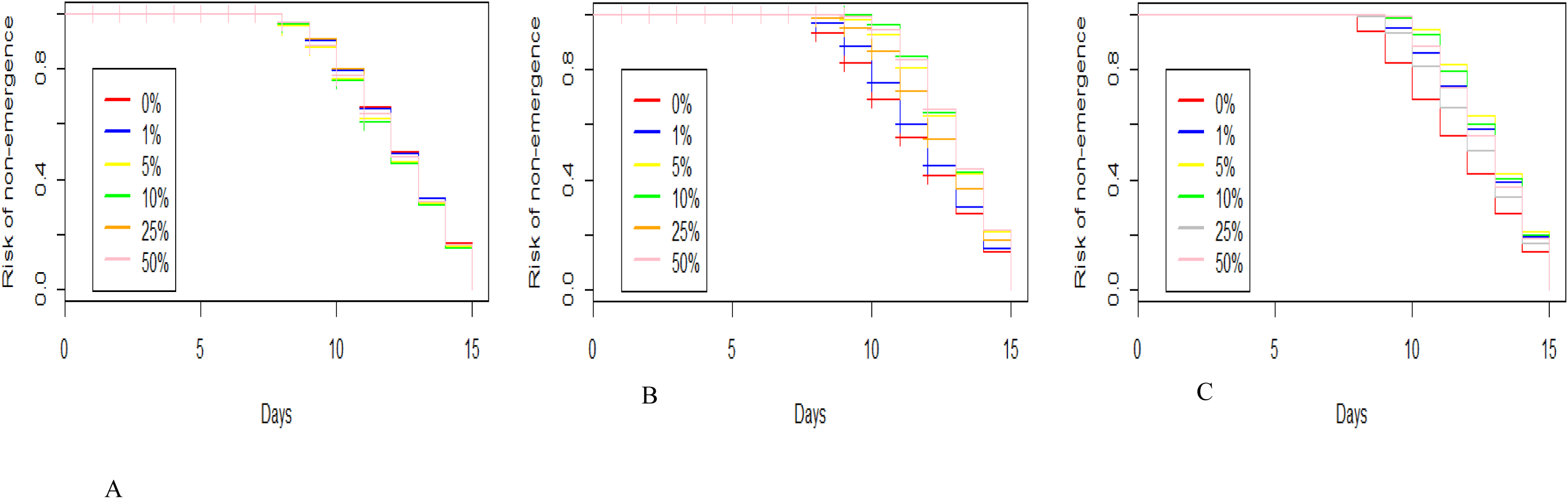
Development time failure measured in days, as the risk to die before adulthood. Eggs were reared on standard *Drosophila* media that was supplemented with differing concentrations of octanoic acid – 0%, 1%, 5%, 10%, 25%, 50%. The graphs are arranged according to species: *D. sechellia* (A), inbred *D. melanogaster* (B) and outbred *D. melanogaster* (C).

When comparing the development times across species, there is little variation in the time taken to develop on different concentrations of octanoic acid in *D. sechellia* (Figure 7). In comparison, inbred *D. melanogaster* take the shortest time to develop when reared at low concentrations, with time increasing as concentration increases. In outbred *D. melanogaster*, flies reared at low concentrations (e.g. 0% and 1%) take the shortest time to develop, similar to inbred *D. melanogaster*. However, outbred *D. melanogaster* flies reared at the higher concentrations (e.g. 25% and 50%) take less time than those reared at middling concentrations, such as 5% and 10%.

#### Female offspring adult weight

In *D. sechellia*, female offspring reared on a 0% acid concentration weighed significantly more than females reared on 1% (T_5_=-4.244; P<0.001), 10% (T_5_=-2.581; P=0.010) or 25% (T_5_=-2.294; P=0.022), but not 5% (T_5_=-1.806; P=0.071) or 50% (T_5_=1.109; P=0.2684) (Figure 8). Females reared on 50% acid concentration were significantly heavier than flies reared on all other concentrations, except 0%.

**Figure 8.**
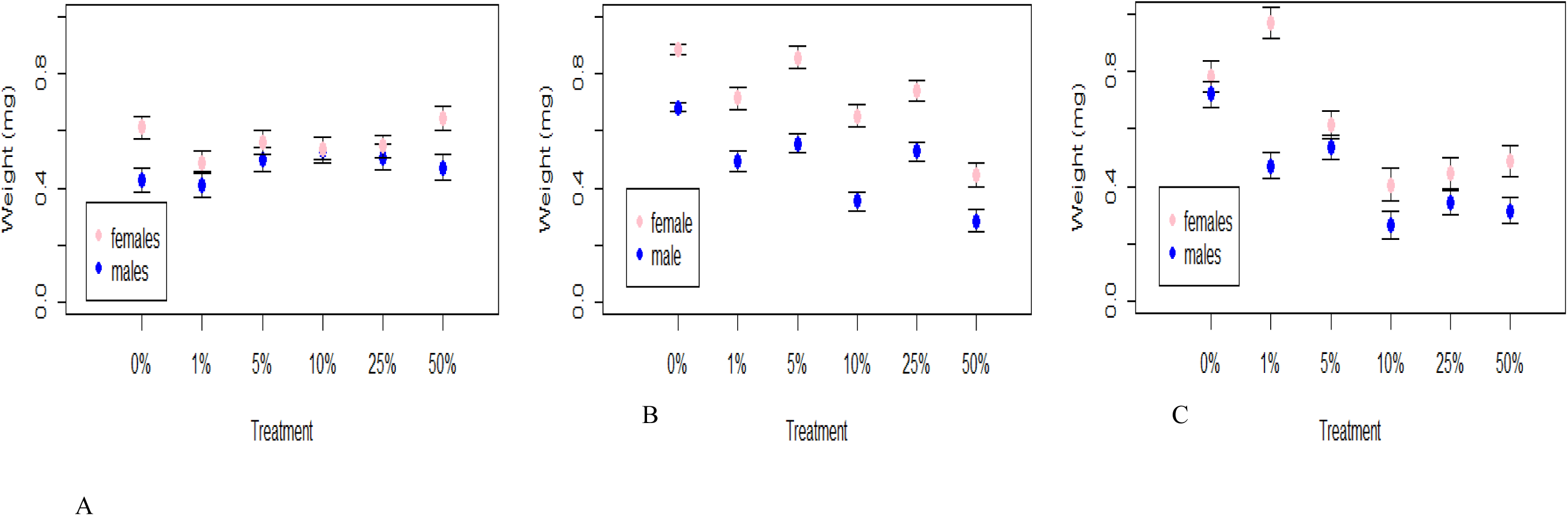
Weight of adult offspring (mg) of newly emerged adults, when their parents were reared on different concentrations of octanoic acid – 0%, 1%, 5%, 10%, 25%, 50%. The graphs are arranged according to species: *D. sechellia* (A), inbred *D. melanogaster* (B) and outbred *D. melanogaster* (C). Females are shown here by the pink dots and males represented by the blue.

In inbred *D. melanogaster* offspring, females reared on 0% acid concentration weighed significantly more than all other treatments, except 5% (T_5_=-1.265; P=0.207): 1% (T_5_=-7.877; P<0.001), 10% (T_5_=-10.339; P<0.001), 25% (T_5_=-6.760; P<0.001) and 50% (T_5_=-19.368; P<0.001). Flies reared on 50% acid concentration weighed significantly less than all other treatments: 1% (T_5_=9.573; P<0.001), 5% (T_5_=14.628; P<0.001), 10% (T_5_=7.163; P<0.001), 25% (T_5_=10.603; P<0.001).

Female offspring of outbred *D. melanogaster* reared on 1% acid concentration weighed significantly more than all other concentrations: 0% (T_5_=-4.853; P<0.001), 5% (T_5_=--9.595; P<0.001), 10% (T_5_=-14.142; P<0.001), 25% (T_5_=-13.217; P<0.001) and 50% (T_5_=-12.149; P<0.001). Females reared on 0% acid concentration weighed significantly more than 5% flies (T_5_=-4.704; P<0.001), 10% (T_5_=-9.727; P<0.001), 25% (T_5_=-8.777; P<0.001) and 50% (T_5_=-7.661; P<0.001).

Female weight has highly variable depending on species (Figure 8). Overall, female *D. sechellia* weighed less than inbred *D. melanogaster* flies, but there was less difference in weight across octanoic acid concentration. Female weight was more variable in both inbred and outbred *D. melanogaster* strains, with flies generally weighing more at low concentrations compared to high.

#### Male offspring adult weight

In male *D. sechellia* offspring, 0% flies weighed significantly less than 5% flies (T_5_=2.346; P=0.19), 10% flies (T_5_=3.401; P<0.001) and 25% flies (T_5_=2.630; P=0.008) (Figure 8). Male offspring reared at 1% concentration weighed significantly less than 5% (T_5_=2.797; P<0.005), 10% (T_5_=3.841; P<0.001) and 25% flies (T_5_=3.081; P=0.002).

In male inbred *D. melanogaster* offspring, unlike female flies, 0% flies weighed significantly more than all other treatments: 1% (T_5_=-9.574; P<0.001), 5% (T_5_=-6.607; P<0.001), 10% (T_5_=-16.772; P<0.001), 25% (T_5_=-7.971; P<0.001) and 50% (T_5_=-19.016; P<0.001). Flies reared on 1% acid concentration weighed significantly more than 10% flies (T_5_=-5.603; P<0.001) and 50% flies (T_5_=-7.993; P<0.001) but weighed significantly less than flies reared on 5% acid concentration (T_5_=2.516; P=0.012).

In comparison, male outbred *D. melanogaster* offspring reared at 0% weighed significantly more than flies reared on 1% acid concentration (T_5_=-7.564; P<0.001), 5% (T_5_=-5.830; P<0.001), 10% (T_5_=-13.483; P<0.001), 25% (T_5_=-11.467; P<0.001) and 50% (T_5_=-12.639; P<0.001). Males reared on 1% and 5% acid concentrations weighed significantly more than flies reared on 10% (T_5_=-6.028; P<0.001, T_5_=-8.137; P<0.001, respectively), 25% (T_5_=-3.826; P<0.001, T_5_=-5.929; P<0.001, respectively) and 50% (T_5_=-4.840; P<0.001, T_5_=-7.022; P<0.001, respectively).

Similar to females, male weight was also highly variable across species (Figure 8). Male *D. sechellia* weight varied less than both inbred and outbred *D. melanogaster* males, with *D. sechellia* males always weighing more than both inbred and outbred *D. melanogaster* males at high concentrations, such as 50%.

#### Bacterial analysis

Bacterial colony growth was observed in all treatments, with both greater diversity and greater abundance of bacteria found in the *D. melanogaster* inbred and outbred flies (Table 3). Sanger sequencing data identifies colony 1 as *Lactobacillus plantarum*; colony 2 as *Paenibacillus sp*. and colony 3 as *Bacillus cereus*. In all the *D. sechellia* adult flies and almost all the outbred *D. melanogaster* flies, only *L. plantarum* growth was observed. This is in comparison to the adult inbred *D. melanogaster* flies, which exhibited substantial growth of both *L. plantarum*, *Paenibacillus sp*. and *B. cereus* bacteria. *Paenibacillus sp*. and *B. cereus* colonies appear to be present in higher numbers when the *D. melanogaster* strains are reared at higher concentrations of the octanoic acid. The number of colonies identified of each bacterial species, from each *Drosophila* species, is included within the appendix (Table 4).

**Table 3.** Presence or absence of bacterial species detected in the midgut of adult *D. sechellia* and both outbred and inbred *D. melanogaster* flies, when reared on diets containing different concentrations of octanoic acid. Presence of a certain bacterial species is denoted with a tick (✓) and absence is with a cross (✗).

| Species | Concentration | *L. plantarum* | *Paenibacillus sp.* | *B. cereus* |
| --- | --- | --- | --- | --- |
| *D. sechellia* | 0% | ✓ | × | × |
| *D. sechellia* | 1% | ✓ | × | × |
| *D. sechellia* | 5% | ✓ | × | × |
| *D. sechellia* | 10% | ✓ | × | × |
| *D. sechellia* | 25% | ✓ | × | × |
| *D. sechellia* | 50% | ✓ | × | × |
| Outbred *D. melanogaster* | 0% | ✓ | × | × |
| Outbred *D. melanogaster* | 1% | ✓ | × | × |
| Outbred *D. melanogaster* | 5% | ✓ | × | × |
| Outbred *D. melanogaster* | 10% | ✓ | × | × |
| Outbred *D. melanogaster* | 25% | ✓ | × | × |
| Outbred *D. melanogaster* | 50% | ✓ | ✓ | × |
| Inbred *D. melanogaster* | 0% | ✓ | × | × |
| Inbred *D. melanogaster* | 1% | ✓ | × | × |
| Inbred *D. melanogaster* | 5% | ✓ | ✓ | ✓ |
| Inbred *D. melanogaster* | 10% | ✓ | ✓ | ✓ |
| Inbred *D. melanogaster* | 25% | ✓ | ✓ | ✓ |
| Inbred *D. melanogaster* | 50% | ✓ | ✓ | ✓ |

#### Octanoic acid aversion trials

Males and females were tested separately to determine if there was a difference in aversion rate according to sex. No difference was observed so subsequent analysis was performed with both sexes grouped together and separated according to species. Unselected *D. sechellia* (hereon known as Dsech ST) were found to prefer the food containing octanoic acid, in comparison to the unselected *D. melanogaster* stock population (hereon known as Dmel ST), which were significantly more averse (F_5_=-2.124; P=0.027). First generation *D. melanogaster* (Dmel 1) flies that had been reared on a diet supplemented with *D. sechellia* gut microbiota, were significantly more averse to the food containing octanoic acid than Dsech ST flies (F_5_=-2.541; P=0.011). There was no difference in aversion of octanoic acid found between Dsech ST and tenth generation *D. melanogaster* (Dmel 10) flies that had been reared on a diet supplemented with *D. sechellia* gut microbiota (F_5_=1.371; P=0.170). Dmel 10 flies were also found to be significantly less averse to octanoic acid than Dmel ST (F_5_=2.889; P=0.003); with no significant difference shown between Dmel ST and Dmel 1 flies (F_5_=-0.973; P=0.330). Notably, Dmel 1 were found to be significantly more averse to the food containing octanoic acid than Dmel 10 flies (F_5_=3.774; P<0.001).

**Table 5.** Proportions of times flies from each treatment group – *D. sechellia, D. melanogaster, D. melanogaster* 1^st^ generation and *D. melanogaster* 10^th^ generation – chose either the diet with octanoic acid present, or the diet without, when placed in the aversion assays.

| Species | Choice |  |
| --- | --- | --- |
|  | Proportion with octanoic acid | Proportion without |
| *D. sechellia* | 0.60 | 0.40 |
| *D. melanogaster* | 0.35 | 0.65 |
| *D. melanogaster* 1st gen | 0.29 | 0.71 |
| *D. melanogaster* 10th gen | 0.69 | 0.31 |

## Discussion

Our results are the first to provide evidence that the gut microbiota may have played a role in host specialisation in *Drosophila sechellia*. We characterised the gut microbiota of wild-type, laboratory reared *D. sechellia* and demonstrated the impact that a changing diet has on gut microbiota. We found that by rearing *D. sechellia* on a fruit diet similar in nutritional properties to its natural host plant but without the toxins, we observe a microbiota of very similar diversity. We then showed the effect that altering the gut microbiota via diet has on subsequent life history traits, larval, pupal and adult weight, with little difference observed between larvae and pupae of all diet types. The only difference was that adults reared on the standard laboratory diet (ASG) weighed significantly less than noni reared flies. Noni flies had significantly higher development failure than salak flies, but they also weighed more at adult emergence, suggesting they have greater fitness.

As predicted, differences in survival ability, development time and resulting offspring adult weight were shown across the three different study groups – *D. sechellia*, inbred *D. melanogaster* and outbred *D. melanogaster –* when exposed to increasing concentrations of toxic, octanoic acid. *D. sechellia* generally exhibited higher survival rates in both males and females, when exposed to higher concentrations of octanoic acid. Inbred and outbred *D. melanogaster* exhibited similar survival abilities, with higher mortality seen at higher acid concentrations. The development time of offspring whose parents were reared on differing concentrations differed according to species, with *D. sechellia* and outbred *D. melanogaster* exhibiting a similar development time, and with more variance seen in inbred *D. melanogaster*. Offspring weight was also more variable in inbred and outbred *D. melanogaster* compared to *D. sechellia*, with both strains of *D. melanogaster* weighing more than *D. sechellia* at lower concentrations, but this effect was reversed at higher concentrations. Similar to previous results (Chandler et al., 2011), when reared on a standard *Drosophila* diet supplemented with octanoic acid, *D. sechellia* gut microbiota was found to be consistent to those isolated from the natural host plant, *M. citrifolia*. The bacteria isolated from *D. sechellia* was characterised as *L. plantarum*. This was also present in the gut of both inbred and outbred *D. melanogaster*, with *B. cereus* and *Paenibacillus sp.* also identified.

Little attention has been paid to the gut microbiota of *D. sechellia*, with the focus instead on genetic adaptation to its toxic host plant. Chandler et al. (2011) characterised the gut bacteria of wild *D. sechellia* found feeding on noni and determined that the natural gut microbiota of this species is dominated by a single *Lactobacillales*. This is in stark contrast to other fruit-feeding, closely related species of *Drosophila* that exhibit considerably greater bacterial diversity, such as wild *D. melanogaster* which host a number of bacterial genera, including *Enterobacteriales*, *Burkholderiales* and *Pseudomonadales*. Here, we show that the gut microbiota of wild-type *D. sechellia* is diverse when individuals are kept under laboratory conditions on a formulated diet. We determined that both males and female guts contain *Paenibacillus sp.* and *Bacillus cereus.* Although evidence has shown that the gut microbiota of laboratory reared species is considerably less diverse than their wild counterparts (Brummel et al., 2004; Roh et al., 2008), some studies do show deviation from the typically found bacterial genera of *Lactobacillus*, *Acetobacters* and *Enterobacter* in laboratory reared flies (e.g. Ren et al., 2007). It could therefore be argued that these genera of bacteria are present in wild populations of *D. sechellia*, but only thrive in great enough numbers for detection when placed onto a diet that encourages their growth. A bacterial pathogen, *Paenibacillus* species are known to be present in honeybee larvae and are responsible for colony collapse by causing American Foulbrood (e.g. Genersch, 2010).

Both *Paenibacillus nanensis* and *B. cereus* have been discovered in wild populations of *Drosophila ananassae* (Maji et al., 2013), although their function or effect on the host is as yet unknown. Presence of these pathogens in both strains on *D. melanogaster* may be due to the natural inability and avoidance of *D. melanogaster* to tolerate the octanoic acid. Although, previous studies have reported death on contact with either the noni or octanoic acid (R’Kha et al., 1991; Dekker et al., 2006), our study shows *D. melanogaster* are able to survive on moderate to high concentrations of the acid for a considerable period. This may be due to the strains of *D. melanogaster* used. Susceptibility to bacterial pathogens may be unwanted side effect of this survival ability, due to a weakened immune system. This may particularly be the case for the inbred strain of *D. melanogaster*, where a lack of genetic diversity may render individuals more susceptible to pathogen colonisation (e.g. Alarco et al., 2004).

When individuals are then transferred onto the natural host noni, the gut microbiota simplifies to a single species - *Lactobacillus plantarum –* as similarly shown in previous studies (Chandler et al., 2011). It could be suggested that colonies of *Lactobacillus plantarum* dominate when individuals are transferred onto noni, due to this bacterium acting as a detoxifying agent by metabolising the toxic compounds present in noni. In humans, *L. plantarum* is responsible for protecting the urogenital and intestinal tracts from infection from pathogenic bacteria (Reid and Burton, 2002). In *Drosophila*, *L. plantarum* has similarly been shown to protect against colonisation of pathogens in the gut (Ryu et al., 2008), by digesting sugars to produce lactic acid, which inhibits the growth of non-commensal organisms and promotes the growth of *Lactobacilli* that thrive in low pH conditions (e.g. Kleerebezem et al., 2003). It is also responsible for promoting larval growth when nutrients are scarce (Storelli et al., 2011), and plays a role in mating preferences (e.g. Sharon et al., 2010). Despite the clear role that *L. plantarum* plays on *D. sechellia* host physiology and likely role in digestion of toxic compounds, high levels of *L. plantarum* were also found when flies were reared on salak. Therefore, the dominance of *L. plantarum* may simply be due to the acidic conditions provided by both fruits. Further work is needed to elucidate the links between these two components.

The weight of individuals at different life stages greatly varied depending on the diet on which they were reared. At the larval stage, no difference in weights was observed across any of the treatments, yet at the pupal stage, ASG pupae weighed significantly more than those reared on noni or salak. As such high abundances of *Lactobacillus plantarum* were found in the gut across all treatments; it could be that *L. plantarum*, which is known to promote larval growth under conditions where nutrients are scarce, is compensating for the host developing on this laboratory formulated diet. In contrast, at adulthood, both male and female ASG reared flies weighed significantly less than both noni and salak flies. One reason for this difference in weights at adulthood may be due to the presence of *B. cereus* and *Paenibacillus sp.* in the adult ASG flies that are not present in individuals reared on noni or salak*. B. cereus* and *Paenibacillus sp.* have been reported in wild populations of *D. ananassae* (Maji et al., 2013), with some studies showing that the immune responses produced by *D. melanogaster* individuals in defence of the pathogen *B. cereus*, can have detrimental effects on life span (Ma et al., 2012; Ma et al., 2013). Therefore, it could be that the immune responses elicited by *D. sechellia* when individuals are reared on the less–preferred diet of ASG override the beneficial effects of *L. plantarum* to negatively affect adult weight.

As predicted, both female and male *D. sechellia* were able to survive on a diet containing the highest concentration of octanoic acid (50%) with around 50% of individuals still alive at the end of the assay. Little differences were observed between any of the different acid concentrations, with the main difference that males reared on diets supplemented with 10% acid displayed a higher survival than those reared on a diet containing 0% octanoic acid. This suggests that addition of octanoic acid to *D. sechellia* is beneficial to survival, at least at lower concentrations. The ability of *D. sechellia* to survive at high concentrations of octanoic acid is somewhat to be expected as *M. citrifolia’s*, main toxic constituent is octanoic acid, although there is some variation in the natural concentrations found, with some studies reporting 58% (Farine et al., 1996) and others 70% (Pino et al., 2010).

Males and females from both the inbred and outbred lines of *D. melanogaster* were able to survive when placed onto a diet containing all concentrations of octanoic acid (Legal et al., 1994; Legal et al., 1999), however survival at high concentrations was lower than *D. sechellia* flies. This result is somewhat surprising as previous studies have noted that *D. melanogaster* dies upon contact with *M. citrifolia*, with most doing so within one hour (Legal et al., 1994; Legal et al., 1999). Survival of both sexes in the inbred *D. melanogaster* strain was significantly reduced at higher concentrations compared to a non-acidic diet, with nearly all individuals recorded dead at the end of the time period. The survival ability of outbred *D. melanogaster* was substantially better than the inbred strain, with only around 50% of females and 30% of males recorded as dead when reared at 50% acid concentration, at the end of the study. The outbred *D. melanogaster* strain may have a better ability to survive on the octanoic acid due to it being a wild-type strain and maintaining genetic diversity. Further, as *D. sechellia* is a sister species of *D. melanogaster*, the outbred or wild-type strain are more likely to share more genetic information than the inbred strain, including genes that underpin resistance to *M. citrifolia*. Animals with increased genetic diversity are also known to adapt to stress better than those with reduced genetic diversity (e.g. Bell, 2013).

The development time of offspring that emerged from adults that were reared on different concentrations of octanoic acid was measured. In *D. sechellia*, no differences were observed in the development time of offspring across any of the octanoic acid concentrations. Similarly, the outbred strain of *D. melanogaster* also exhibits little variation in development time between all octanoic acid concentrations, but flies reared at 0% took the shortest time to develop. This is in comparison to the inbred *D. melanogaster*, in which flies reared at the 0% acid concentration took significantly longer to develop than flies reared at higher concentrations. It is surprising that outbred *D. melanogaster* flies displayed a similar development time to *D. sechellia*. This is likely due to the conserved genetic diversity in the outbred line. Similar studies have shown that inbred *D. melanogaster* lines have a significantly reduced lifespan when exposed to dietary-restrictions, compared to that of outbred lines such as Dahomey (Grandison et al., 2009).

No difference was observed in adult weight of *D. sechellia* offspring, in parent flies reared on both 0% and 50% octanoic acid concentrations, with little variation observed in both males and females overall. In comparison, outbred *D. melanogaster* flies reared at higher concentrations, between 10% and 50%, exhibited a significantly reduced weight in both sexes, with flies reared at 1% in females, and 0% in males weighing the most. Similar to the outbred line, male and female inbred *D. melanogaster* had a more variable weight range, with flies reared at 0% acid concentration weighing significantly more than flies reared at higher concentrations. As the concentration of octanoic acid in noni is between 58% (Farine et al., 1996) and 70% (Pino et al., 2010), the results for *D. sechellia* are contrary to expected. It could be predicted that *D. sechellia* reared at 0% concentration of octanoic acid would weigh less as this does not mimic its natural host plant. However, as this strain, although they are outbred, have been maintained in the laboratory since 1980, they may have become better adapted to the laboratory diet over time (e.g. Telonis-Scott et al., 2006). Variation between the weights of inbred and outbred strain flies is likely due to the conserved genetic diversity in the outbred line. Similar studies have shown that inbred *D. melanogaster* lines have a significantly reduced lifespan when exposed to dietary-restrictions, then that of outbred lines such as Dahomey (Grandison et al., 2009).

*Drosophila melanogaster* is known to die upon contact with the natural host plant of its sister species, *D. sechellia*, and as such has evolved mechanisms to detect and avoid this fruit (Legal et al., 1994; Legal et al., 1999). Using a series of aversion assays, we highlighted the differences in behavioural response to the presence of octanoic acid – the main chemical component of noni and the chemical responsible for its pungent scent and toxic nature. Similar to previous results, stock population *D. melanogaster* were shown to be significantly more averse to octanoic acid than *D. sechellia* (Legal et al., 1999; Dekker et al., 2006). This is undoubtedly due to the presence of the OBPs (OBP57d and OBP57e) present in both sexes of *D. sechellia* that attract the flies to the octanoic acid within the fruit. Detection of the octanoic acid scent increases oviposition in females and so females seek the fruit in order to lay (Legal et al., 1999).

A significant difference was found between *D. sechellia* and first-generation *D. melanogaster* flies that had been supplemented with *D. sechellia* gut microbiota. Here, first-generation *D. melanogaster* flies were significantly more averse to the food containing octanoic acid, and thus resembled the behaviour of standard stock population *D. melanogaster* (Legal et al., 1999; Dekker et al., 2006). However, no difference was found in preference for tenth-generation *D. melanogaster* flies compared with *D. sechellia*, showing that the aversion response was significantly reduced between first-generation and tenth-generation *D. melanogaster* individuals. This result potentially illustrates the first step in an organism specialising to a novel and toxic host. The genetic basis for aversion in *D. melanogaster* is due to the absence of the odorant-binding proteins (OBPs) OBP57d and OBP57e that are present on the gustatory sensilla on the legs of *D. sechellia* (e.g. Amlou et al., 1998; Jones, 2005; Matsuo et al., 2007; Dworkin and Jones, 2009). These enable the flies to detect the odour from up to 150m away (R’Kha et al., 1991). However, no attention has been paid to the role of the gut microbiota in evolutionary preference and ability to synthesis toxic octanoic acid. We suggest that the gut microbiota can interact with the genetic mechanisms within the fly to override the natural aversion response, and thus contribute to the role of specialisation in this insect.

The gut microbiota of *D. melanogaster* is diverse but highly dependent on a number of factors, including diet (e.g. Sharon et al., 2010), age (Zerofsky et al., 2005), or strain (e.g. discussed in Heys et al., 2018b). When reared on a standard *Drosophila* diet (0% acid concentration), our study determined that *L. plantarum* is present within both the *D. melanogaster* and *D. sechellia* gut. The results of the experimental evolution line of *D. melanogaster* supplemented with *D. sechellia* gut microbiota that we created, supports our argument that *L. plantarum* acts a detoxifying agent within the *Drosophila* gut. Octanoic acid is responsible for the majority of toxicity within the fruit (Farine et al., 1996; Jones, 1998). We predict the gut is able to withstand the toxicity of both the octanoic and hexanoic acids, via metabolising the toxic compounds into less harmful products which are able to be digested. Previous studies have shown that pH can determine the gut microbial composition (see Overend et al., 2016) and it could therefore be argued that sole presence of *L. plantarum* within the *D. sechellia* gut is due to pH alone. By determining the difference found between the first and tenth generation experimental evolution lines in aversion to octanoic acid, we can dispute the idea that the high colony numbers of *L. plantarum* is simply due to the increased pH. For example, similar scenarios can be viewed in the mealworm, *Tenebrio molitor*, which detoxifies the cell walls of fungi and bacteria within its diet (Genta et al., 2006), and the coffee berry borer, *Hypothenemus hampei*, a specialist of coffee plants where *Pseudomonas* species within the gut microbiota metabolise caffeine - a toxic alkaloid (Ceja-Navarro et al., 2015).

This is the first time the gut microbiota of *D. sechellia* has been examined in laboratory conditions under different dietary treatments. Our results are in-keeping with others that characterise the microbiota of wild-caught *D. sechellia* (Chandler et al., 2011) despite the fact that our population has been reared in the laboratory on a formulated diet for a number of years. Here we demonstrate that when *D. sechellia* are reared on its natural host at any time point, a shift in the gut microbiota can be seen. Our results are the first to show the direct change in gut microbiota when the same individuals are moved between vastly different diets – the natural host plant and a laboratory diet. Although we determine these differences in microbiota across dietary treatments, further work is needed to disentangle the effect of pH on the gut microbiota, from a shift in microbiota that enables specialisation within this species. The present study provides evidence that the gut microbiota is responsible for specialisation in *D. sechellia*. *D. melanogaster* is known to be highly averse to the scent profile of octanoic acid, in comparison to *D. sechellia* in which it is a chemoattractant (Louis and David, 1986; R’Kha et al., 1991; Higa and Fuyama, 1993; Amlou et al., 1998; Legal et al., 1999). By creating experimental evolution lines of outbred *D. melanogaster* that are supplemented with *D. sechellia* gut microbiota, we have significantly reduced aversion of *D. melanogaster* to octanoic acid after only ten generations. In particular we suggest that *Lactobacillus plantarum*, the main bacterial constituent of the *D. sechellia* gut, acts as a detoxifying agent to metabolise the toxic octanoic acid compound – the main chemical present in the natural host plant, M. *citrifolia*. Little is known as to the origins of speciation of *D. sechellia* from its sister species, however, our results suggest that shifts in the gut microbiota may have led to ecological divergence, and later speciation, within this species. We have demonstrated an evolutionary shift in preference to food containing octanoic acid in *D. melanogaster*, to which it is naturally averse. Reducing aversion to a novel host plant could be the first step in ecological and evolutionary divergence. Further work is needed to understand how *L. plantarum* metabolises octanoic acid into presumably harmless components that can be utilised by the host.

## Acknowledgements

This work was supported by the Natural Environment Research Council (grant number NE/L002450/1). The authors wish to thank Andrea Betancourt, Alistair Darby, Greg Hurst, and Tom Price, discussions with whom greatly improved this study.

## Appendix

**Table 1.** Sample sizes for the measurements of development time for each species and strain measured: *D. sechellia*, *D. melanogaster* inbred, *D. melanogaster* outbred.

| Species | Treatment | Sample size |
| --- | --- | --- |
| Inbred *D. melanogaster* | 0% | 279 |
| Inbred *D. melanogaster* | 1% | 275 |
| Inbred *D. melanogaster* | 5% | 202 |
| Inbred *D. melanogaster* | 10% | 157 |
| Inbred *D. melanogaster* | 25% | 117 |
| Inbred *D. melanogaster* | 50% | 111 |
| Outbred *D. melanogaster* | 0% | 235 |
| Outbred *D. melanogaster* | 1% | 213 |
| Outbred *D. melanogaster* | 5% | 185 |
| Outbred *D. melanogaster* | 10% | 124 |
| Outbred *D. melanogaster* | 25% | 121 |
| Outbred *D. melanogaster* | 50% | 60 |
| *D. sechellia* | 0% | 152 |
| *D. sechellia* | 1% | 139 |
| *D. sechellia* | 5% | 198 |
| *D. sechellia* | 10% | 136 |
| *D. sechellia* | 25% | 174 |
| *D. sechellia* | 50% | 176 |

**Table 2.**
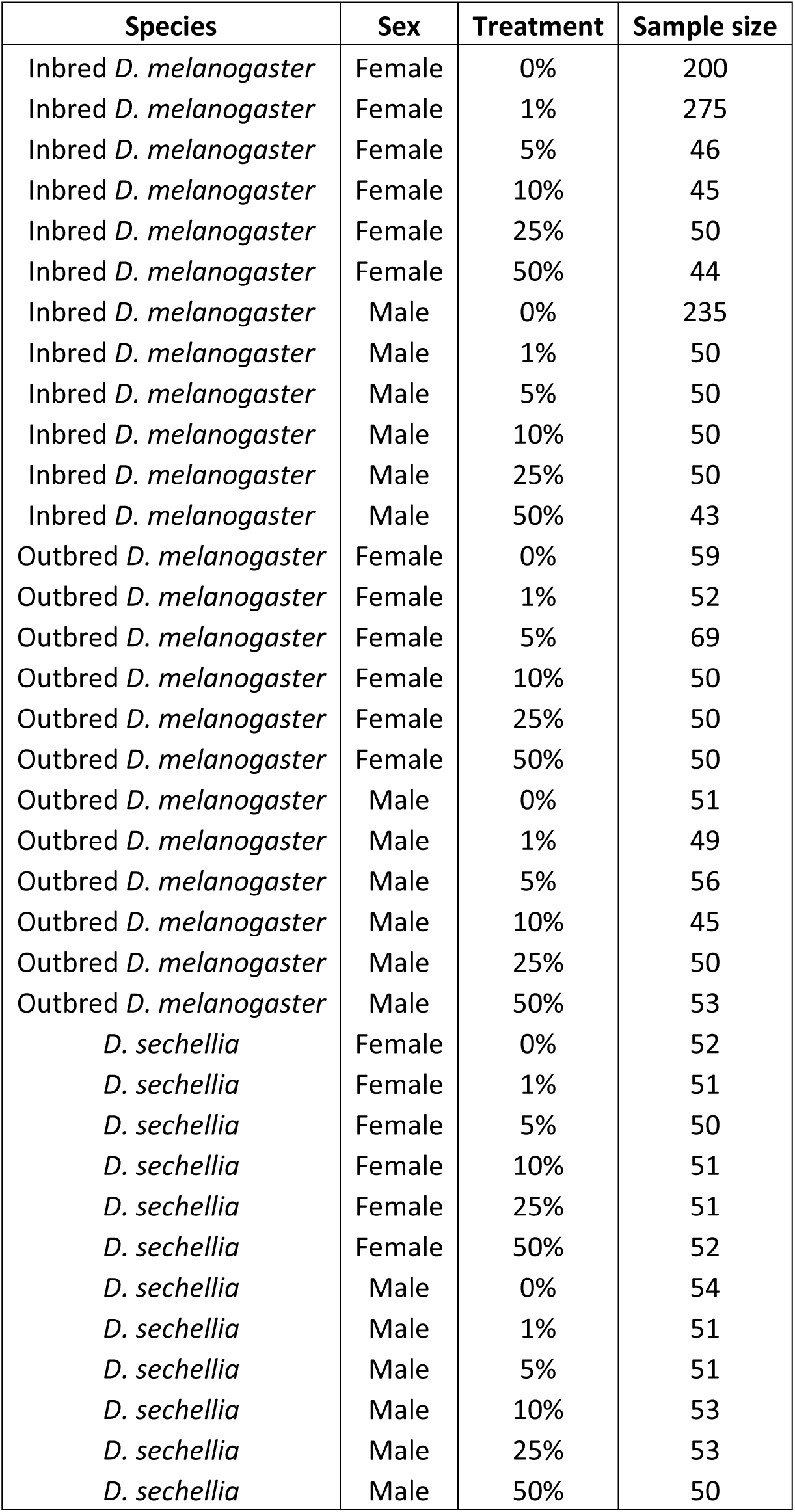
Sample sizes for the measurements of adult emergence weight for each species, sex and strain measured: *D. sechellia*, *D. melanogaster* inbred, *D. melanogaster* outbred.

